# *Phytophthora* root rot induces compositional and functional changes in avocado rhizosphere bacterial communities

**DOI:** 10.1101/2025.11.20.689515

**Authors:** Rosaura G. Alfaro-García, Pablo Vargas-Mejía, Violeta Patiño-Conde, Eria A. Rebollar, José A. Guerrero-Analco, Julio Vega-Arreguín, Frédérique Reverchon, Alfonso Méndez-Bravo

**Affiliations:** Red de Diversidad Biológica del Occidente Mexicano, Instituto de Ecología, A. C., Centro Regional del Bajío. Av. Lázaro Cárdenas 253, Centro. 61600 Pátzcuaro, México; Laboratorio Nacional de Análisis y Síntesis Ecológica, Escuela Nacional de Estudios Superiores, Unidad Morelia, Universidad Nacional Autónoma de México, Antigua carretera a Pátzcuaro 8701, Ex Hacienda de San José de la Huerta. 58190 Morelia, México; Laboratorio de Ciencias Agrogenómicas and Laboratorio Nacional PlanTECC, Universidad Nacional Autónoma de México. Blvd. UNAM 2011. 37684 León, México; Centro de Ciencias Genómicas, Universidad Nacional Autónoma de México. Av. Universidad s/n, Universidad Autónoma del Estado de Morelos. 62210 Cuernavaca, México; Red de Estudios Moleculares Avanzados, Instituto de Ecología, A.C. Carretera Antigua a Coatepec 351, El Haya. 91073 Xalapa, México; SECIHTI, Ciudad de México, México

**Author notes:** Co-corresponding authors: Frédérique Reverchon; Alfonso Méndez-Bravo.

**Keywords:** Cry-for-help, Dysbiosis, Metatranscriptomics, *Persea americana*, *Phytophthora cinnamomi*, Rhizobacteria

## Abstract

Understanding how plant pathogens modulate the rhizosphere microbiota is essential to integrated disease management. Here, we assessed the compositional and functional shifts in the avocado rhizosphere bacteriome induced by *Phytophthora cinnamomi* to elucidate the microbial functions modulated by the infection and identify taxa potentially recruited by the plant as a defense response. Through metabarcoding with metatranscriptomics, we showed that *Phytophthora* root rot (PRR) induced compositional shifts in bacterial communities, leading to the enrichment of members of MND1, RB41 and *Nitrospira*. Functional analysis showed that this enrichment may be due to the release of nutrients following root rot, as carbohydrate metabolism was stimulated in rhizobacterial communities of infected trees. We detected evidence of a cry-for-help strategy by the infected plant, as the most active genera in the rhizosphere of PRR-symptomatic trees up-regulated genes associated with stress response and cell signaling, suggesting that they were recruited to mitigate the adverse effects of infection. Our findings highlight the need to combine compositional and functional microbiome data to differentiate between taxa attracted by nutrient release and those actively recruited by the plant. The interactions of the latter with the pathogen should be further studied, as they may constitute promising biocontrol agents.

## Introduction

The rhizosphere microbiota is critical for plant health and productivity (Trivedi et al., 2020). By improving plant nutrient uptake, producing phytohormone-like compounds, stimulating their host plant defense response and increasing its tolerance to abiotic and biotic stressors, rhizosphere microbial communities contribute to their host’s growth and fitness (Compant et al., 2025; Méndez-Bravo et al., 2018). Research from the last two decades have shed light on the factors driving the assembly of rhizosphere microbial communities, which are both host-dependent (e.g., plant genotype or developmental stage; Chaparro et al., 2013; Mendes et al., 2018; Philippot et al., 2013) and host-independent (e.g., soil parameters, climatic variables, or human management practices; Carrasco-Espinosa et al., 2022; Hartmann et al., 2014; Santoyo, 2022). In particular, plant health status has been shown to be a strong determinant of rhizosphere microbial assemblages, as plant infection by pathogens can induce shifts in diversity, composition and function in plant-associated microbiota, leading to a possible dysbiotic state (Alfaro-García et al., 2025; Berendsen et al., 2012). In light of the plant holobiont theory (Vandenkoornhuyse et al. 2015), understanding how plant pathogens modulate the rhizosphere microbiota may hold the key to more efficient strategies for integral disease management, which is particularly relevant in monoculture cropping systems where soil-borne pathogen loads may be accrued (Dou et al., 2025; Hiddink et al., 2010).

*Phytophthora cinnamomi* Rands., a soil-borne oomycete, is considered one of the most devastating pathogens worldwide (Hardham and Blackman, 2018). Root rot caused by *P. cinnamomi* causes severe ecological and economic damages in a wide range of natural and agro-ecosystems (Shands et al., 2025). In avocado (*Persea americana* Mill.), *Phytophthora* root rot (PRR) represents the major limiting factor of the fruit global production, including Mexico, the world’s leading producer and exporter of avocado (Fernández-Pavía et al., 2013; Guevara-Avendaño et al., 2024). The shifts in the avocado rhizosphere microbiome induced by *P. cinnamomi* have been studied at the compositional and structural level, with contrasting results. Yang et al. (2001), using denaturing gradient gel electrophoresis (DGGE), detected that the rhizosphere bacterial community of avocado was more diverse in roots infected with *P. cinnamomi* than in uninfected roots, although uninfected roots were colonized by bacterial genera such as *Pseudomonas*, *Polyangium* and *Cytophaga*, which have been reported as controllers of pathogens (Nana et al., 2023; Yuan et al., 2022). Using shotgun metagenomics, Shu et al. (2019) reported that the avocado rhizosphere microbiota was modified by PRR, and showed an increase in the relative abundance of the ten most representative bacterial taxa in infected plants compared with uninfected trees. More recently, Solís-García et al. (2021) used a metabarcoding approach to report shifts in the composition of avocado bacterial and fungal communities due to PRR, with an increase in the relative abundance of copiotrophic taxa such as Pseudomonadales and Burkholderiales and a concomitant decrease of Actinobacteria and Rhizobiales in the rhizosphere of infected trees. Considering the heterogeneity of the described shifts in avocado rhizobacterial communities driven by PRR, further research should focus on the functional consequences of the disease, to elucidate the microbial processes enhanced or hindered by the infection and identify potential contributions of the rhizosphere microbiome in the plant defense response (Alfaro-García et al., 2025; Flores-Nunez & Stukenbrock, 2024).

In this study, we assessed the impact of PRR on the diversity, composition and gene expression profile of avocado rhizosphere bacterial communities in an orchard located in the world’s leading producing region. We combined a metabarcoding approach with a metatranscriptomic analysis of the rhizosphere bacteriome in asymptomatic and PRR-symptomatic avocado trees to identify potential bacterial taxa and functional markers associated with the disease or contributing to its mitigation.

## Materials and methods

### Study site and rhizosphere soil sampling

The study orchard was located in Peribán, Michoacán (19’ 31’16” N, 102’ 24’54” W) at an altitude of 1,640 m.a.s.l. The mean annual temperature is 19.8 ‘C, with an annual precipitation of 1,445 mm mostly falling between June and September (CONAGUA, 2023). The selected orchard is under organic management and the presence of *P. cinnamomi* has been recorded in recent years. On April 2023, we randomly selected 10 avocado trees that showed symptoms of PRR according to Solís-García et al. (2021) —with defoliation, dead or chlorotic leaves, coupled with small, brittle and/or necrotic roots)— and 10 asymptomatic avocado trees, all aged between eight and 15 years old. The presence of *P. cinnamomi* in the roots of PRR-symptomatic trees was confirmed by culture-dependent methods. Around each selected tree, we drew a circumference no greater than 100 cm and collected the rhizosphere soil adhered to fine roots at four points around the trunk by digging with an ethanol-disinfected shovel (10 cm depth). The four samples were mixed and homogenized to obtain one rhizosphere soil sample per tree, and subsequently divided into two centrifuge tubes (50 ml), one for DNA extraction and the other for RNA extraction. Thirty milliliters of Qiagen’s LifeGuard® Soil Preservation Solution were added to the soil samples aimed for RNA extraction. All rhizosphere soil samples were kept cold with frozen gel bags until the arrival at the lab, where they were stored at -80 ’C until DNA and RNA extraction.

### Soil chemical characterization

Three randomly selected trees per condition (asymptomatic and PRR-symptomatic) were chosen to collect 500 g of soil for chemical characterization. Soil pH was measured in a 1:2 ratio in deionized water (AS-02-NOM-021-RECNAT-2000, SEMARNAT, 2002). Electrical conductivity was registered in a 1:5 soil/water ratio (FAO, 2021b). Available phosphorus (P) was determined with the Olsen method (FAO, 2021a). Total carbon (C) and nitrogen (N) were quantified using the combustion method in an elemental analyzer (LECO, TRuspec model). The potassium chloride extraction method was used to determine nitrate (NO_3_^-^) and ammonium (NH_4_^+^) concentrations (Bremner, 1965). All soil analyses were performed at the Laboratory for Soil Analyses of the *Instituto de Ecología, A.C*., Xalapa, México. A Student’s t-test was used to compare soil chemical characteristics between asymptomatic and PRR-symptomatic avocado trees. There were no significant differences in soil characteristics between samples from the rhizosphere of asymptomatic and PRR-symptomatic avocado trees (Table 1), except for NH_4_^+^, which was higher in the rhizosphere of asymptomatic trees than in that of PRR-symptomatic trees.

**Table 1.** Chemical characteristics (means ± standard deviation, n=3) of rhizosphere soil from asymptomatic and PRR-symptomatic avocado trees.

| Condition | pH | EC | Available P | NH<sub>4</sub><sup>+</sup> | NO<sub>3</sub><sup>-</sup> | Total N | Total C | C/N |
| --- | --- | --- | --- | --- | --- | --- | --- | --- |
|  | 1:2 H<sub>2</sub>O | mS/cm | mg/kg |  |  | % |  | ratio |
| Asymptomatic | 6.49±<br>0.29 | 0.53±<br>0.37 | 178.25±<br>89.30 | 18.04±<br>2.07 | 30.90±<br>10.72 | 0.27±<br>0.11 | 3.60±<br>0.98 | 13.74±<br>1.74 |
| Symptomatic | 6.82±<br>0.14 | 0.26±<br>0.13 | 178.42±<br>85.72 | 12.97±<br>1.02 | 24.92±<br>7.67 | 0.30±<br>0.06 | 3.97±<br>0.96 | 13.38±<br>1.80 |
| Student's t-test | t=-1.74,<br>df=2.83,<br>p = 0.18 | t=-1.22,<br>df=2.46,<br>p = 0.32 | t=-0.002,<br>df=3.99,<br>p = 0.99 | **t=3.80,**<br>**df=2.92,**<br>**p = 0.03\*** | t=0.77,<br>df=3.62,<br>p = 0.47 | t=-0.38,<br>df=3.10,<br>p = 0.72 | t=-0.46,<br>df=3.99,<br>p = 0.66 | t=0.25,<br>df=3.99,<br>p = 0.81 |

### DNA extraction and construction of the 16S rDNA sequencing libraries

DNA extractions from rhizosphere soil were performed with the DNeasy® PowerSoil® Kit (Qiagen) following manufactureŕs instructions. Purity and concentration of extracted DNA were verified using a BioSpectrometer® (Eppendorf). Twenty libraries were constructed for taxonomic profiling of the bacterial communities following the workflow plan outlined in 16S Metagenomic Sequencing Library Preparation of Illumina (2013) with slight modifications. The primers 341F (5‘-CCTACGGGNGGCWGCAG-3’) and 805R (5‘-GACTACHVGGGTATCTAATCC-3’) were used to amplify the V3-V4 region of 16s rRNA to create ∼460 bp amplicons (Illumina, 2013). These first amplicons were dual labeled using the Nextera XT Index kit^TM^ (Illumina). Both PCR reactions (50 µl) were performed following the manufacturer’s instructions for the Multiplex PCR (Qiagen), using a total of 30-40 ng of DNA per amplicon reaction and a total 15 ng per index reaction. The PCR cycling parameters for amplicons and index PCR followed the Illumina (2013) protocol but included one enzyme activation step of 15 minutes at 95°C. Each PCR product was purified with ProNex® Size Selective Purification System following manufactureŕs instructions. The libraries were sequenced paired-end (PE) with a read length of 2 × 300 bp using the Illumina MiSeq^TM^ system at CD Genomics, obtaining an average of 103,646 PE reads per sample.

### RNA extraction

Duplicate RNA extractions were carried out from all rhizosphere soil samples using the RNAeasy® PowerSoil® Total RNA Kit by Qiagen. Prior to the first step of the extraction protocol, each sample and its duplicate were centrifuged at 5,000 g for 5 min to remove the LifeGuard® Soil Preservation Solution as recommended by the manufacturer. The integrity of the extracted RNA was evaluated by the presence of ribosomal RNA (rRNA) bands by electrophoresis. RNA concentration and purity were measured on an Eppendorf BioSpectrometer®. Nine samples with intact RNA from asymptomatic trees were randomly distributed into three equimolar pools and the 10 samples from PRR-symptomatic trees were also distributed into three equimolar pools. Six libraries were then built from the pools at Macrogen, Inc. All the pools received a DNase pre-treatment and, for construction of metatranscriptomic libraries, a NEBNext® rRNA Depletion Kit (Bacteria) was used for rRNA depletion. All libraries were sequenced PE with a read length of 2 × 100 bp using the Illumina NovaSeq 6000^TM^ system at Macrogen, Inc., obtaining an average of 99,056,583 PE reads per sample.

### Bioinformatic and statistical analysis for 16S rRNA sequences

#### Quality control and 16S rRNA amplicon analysis

DNA (16S rRNA) sequences were subjected to a pre-processing analysis to verify sequence quality using the FastQC software v.0.12.1. (Andrews, 2010). The Trimmomatic software v.0.39 was used for removing universal Illumina adapters and low-quality reads (Bolger et al., 2014). The following parameters were used as filters: sequence quality > 30, minimum sequence length > 180 bp and max sequence length 240 bp. Reads were trimmed by 10 bases at the 5’-end and 3’-end. PANDAseq was used to merge PE reads of 16S rRNA gene (Masella et al., 2012).

The composition and structure of the rhizobacterial community was analyzed using the DADA2 pipeline in R v.4.3.3 (Callahan et al., 2016; R Core Team, 2023). After merging, amplicons were trimmed again to 450 bp as minimum sequence length (filterAndTrim function, truncLen argument) and sequences with undetermined bases were removed (filterAndTrim function, maxN argument). Afterwards, sequences were de-replicated to reduce redundancy. An error model was generated by learning the specific error-signature for each sequence. After quality processing, sequences were grouped into Amplicon Sequence Variants (ASVs). Chimeras were removed before taxonomic assignment (removeBimeraDenovo function). Taxonomic assignment was performed using the SILVA v.138.1 database. Taxa with low prevalence (threshold < 5%) were excluded. Singletons, doubletons, mitochondria, chloroplast and eukaryotic sequences were removed before diversity and statistical analyses.

#### Diversity and statistical analyses

The ASV table was rarefied to 36,227 reads using the rarefy_even_depth function implemented in the *Phyloseq* package v.1.46.0 (McMurdie & Holmes, 2013). To calculate the alpha diversity indexes, the estimate_richness function of *Phyloseq* was used. Differences in Observed ASVs, Shannon and Simpson metrics between conditions were tested with the Mann-Whitney-Wilcoxon test using the wilcox.test function, indicating False in the paired argument. Venn diagrams were plotted using the *ggven* package in R. For beta diversity analyses, the Bray-Curtis dissimilarity matrix was calculated from the normalized ASV table and used to build a non-metric multidimensional scaling (NMDS) using the ordinate and plot_ordination functions of *Phyloseq* package, respectively. A permutational multivariate analysis of variance (PERMANOVA) was used to compare the structure of the bacterial rhizosphere community of asymptomatic and PRR-symptomatic avocado trees, using the *vegan* package with 999 permutations. To distinguish true biological differences from potential dispersion effects, we conducted a permutational analysis of multivariate dispersions (BETADISP) using the betadisper and permutest functions with 999 permutations in R. The *DESeq2* package v.1.42.1 was used to compare the differential abundance of bacterial taxa between PRR-symptomatic and asymptomatic trees; significant differences were supported by a Wald test. All statistical analyses were performed using customized scripts with the free software R v.4.3.3. and were considered significant at *P* < 0.05. The analyses were visualized using *ggplot2* v.4.0.0 in R. The 16S rRNA amplicon libraries are available at the Sequence Read Archive of the NCBI under BioProject PRJNA1363127.

### Bioinformatic analysis of metatranscriptomic sequences

#### Quality control and preprocessing

Raw sequence libraries were quality checked using FastQC. Adaptor trimming and removal of low-quality bases were performed with BBDuk from the BBMap suite v.39.06 using the parameters forcetrimleft = 13 and trimq = 10 to ensure high-confidence base calls (https://sourceforge.net/projects/bbmap/). Paired-end reads were subsequently merged using PEAR v0.9.8 with default settings (Zhang, Kobert, Flouri, & Stamatakis, 2013). To eliminate ribosomal RNA, SortMeRNA v.4.3.6 (Kopylova, Noé, & Touzet, 2012) was employed using the SILVA rRNA databases for bacterial and archaeal 16S/23S, and eukaryotic 18S/28S subunits. The --num_alignments 1 parameter was used to retain only the best alignment per read. Reads identified as rRNA were removed for downstream analyses.

#### Read mapping, functional annotation and quantification

Filtered reads were mapped to the NCBI RefSeq bacterial protein database (Tatusova et al., 2014) using DIAMOND v.2.1.6 (Buchfink, Xie, & Huson, 2015) in blastx mode. Only the best-scoring alignment per read was retained to avoid redundant annotations. Alignment results were processed with the SAMSA2 pipeline (Westreich et al., 2018), which extracted protein-level matches and generated count matrices summarizing gene-, taxon- and function-level abundance for all metatranscriptomic libraries.

#### Differential expression and functional enrichment analysis

The NCBI RefSeq bacterial protein database, used for read mapping with DIAMOND, was functionally annotated prior to analysis using EggNOG-mapper v.2 (Cantalapiedra et al., 2021) with the eggNOG v.5.0 database. This provided orthology assignments, functional descriptions, KEGG Orthology (KO) identifiers, and Gene Ontology (GO) terms for all reference proteins, enabling the functional interpretation of mapped reads. Read counts per gene were compiled from the SAMSA2 output and imported into *DESeq2* (Love et al., 2014) for differential expression analysis between asymptomatic and PRR-symptomatic avocado trees. Statistical significance was determined using the Wald test and adjusted for multiple testing using the Benjamini-Hochberg method. Functional enrichment analysis was performed using the *ClusterProfiler* R package (Yu et al., 2012), based on the GO and KEGG annotations assigned to the matched proteins in the annotated reference database. Metatranscriptomic libraries are available at the Sequence Read Archive of the NCBI under BioProject PRJNA1363127.

## Results

### Composition of the rhizosphere bacterial community in avocado trees

A total of 1,969,273 paired reads were obtained from 20 rhizosphere soil samples. After quality filtering, a total of 906,590 reads were retained, representing 25,869 bacterial ASVs. Rarefaction curves showed that rhizosphere samples from both asymptomatic and PRR-symptomatic avocado trees reached a plateau, indicating that most bacterial ASVs were detected (Supplementary Fig. 1).

The PRR infection did not induce significant differences in alpha diversity metrics (Observed richness, Shannon and Simpson) of the rhizosphere bacterial communities compared to asymptomatic trees (Figure 1A). Similarly, PRR did not affect the bacterial community structure in the avocado rhizosphere, as no significant differences were detected between samples from asymptomatic and PRR-symptomatic avocado trees (PERMANOVA df = 1, F = 2.88, *P* = 0.103; Figure 1B). Interestingly, bacterial communities from PRR-symptomatic trees showed more dispersion than those associated with asymptomatic trees. Permutational analysis of multivariate dispersions (BETADISP, *P* = 0.09) confirmed that the observed dispersion could be driven by biological variation (*i.e*., presence or absence of the pathogen) rather than by differences in within-group dispersion.

**Figure 1.**
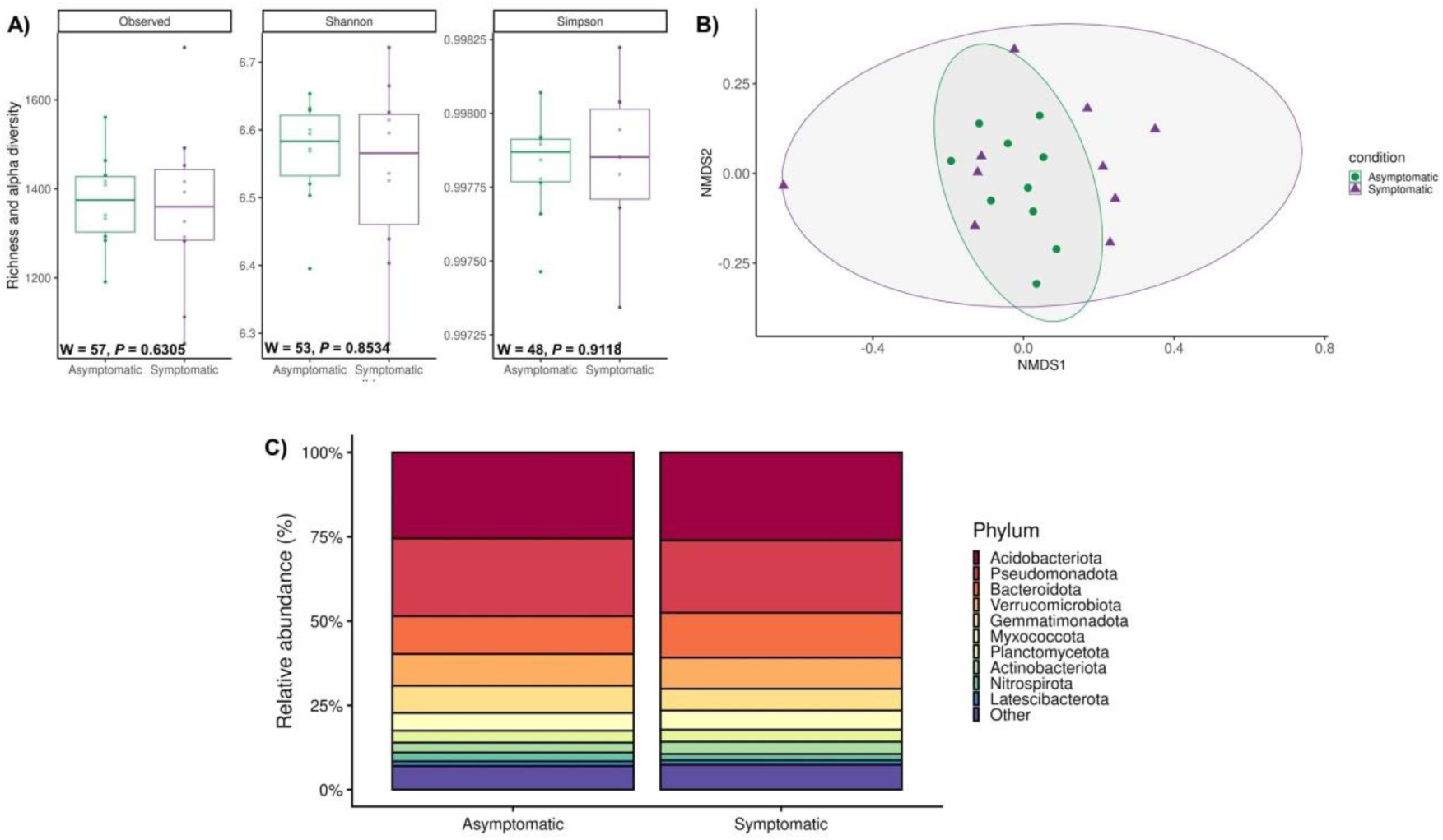
Diversity and composition of the bacterial community in the rhizosphere of asymptomatic and PRR-symptomatic avocado trees. A) Observed richness, Shannon and Simpson indices of rhizobacterial communities associated with asymptomatic and PRR-symptomatic avocado trees. W and *P* values were calculated with the Mann-Whitney-Wilcoxon test. B) Non-metric multidimensional scaling (NMDS) plot based on Bray-Curtis distance of the bacterial community associated to asymptomatic and PRR-symptomatic avocado trees (stress value = 0.19). C) Relative abundance (%) of bacterial phyla associated with asymptomatic and PRR-symptomatic avocado trees. The others category represents phyla with a relative abundance < 1 %.

At the compositional level, the five most abundant bacterial phyla were common to asymptomatic and PRR-symptomatic trees and were Acidobacteriota, Pseudomonadota, Bacteroidota, Verrucomicrobiota and Gemmatimonadota, which together accounted for more than 70 % of the total bacterial community. No significant differences were found in the relative abundance of the main bacterial phyla between conditions (Figure 1C). However, at the genus level, compositional differences were observed between rhizosphere bacterial communities associated with asymptomatic and PRR-symptomatic trees.

From the 482 identified bacterial genera, 303 (62.9 %) were shared by both conditions, 74 (15.4 %) were only found in the rhizosphere of asymptomatic avocado trees and 105 (21.8 %) were exclusively detected in the rhizosphere of PRR-symptomatic trees (Figure 2A). At the ASV level, only 865 (3.8 %) from the 25,080 assigned were shared between the asymptomatic and PRR-symptomatic trees (Figure 2B). Differential abundances between conditions showed that there were more bacterial genera significantly enriched in asymptomatic than in PRR-symptomatic avocado trees (Figure 2C). The *DeSeq* analysis supported by Wald test (*P* < 0.05) showed that rhizosphere bacterial ASVs affiliated with *Rhodanobacter*, *Chryoseolinea*, *Candidatus* Udaeobacter, *Anaeromyxobacter*, *Variovorax*, MND1, RB41 and *Nitrospira* were enriched in asymptomatic avocado trees. In contrast, other ASVs from MND1, RB41 and *Nitrospira* were significantly more abundant in the rhizosphere of PRR-symptomatic avocado trees.

**Figure 2.**
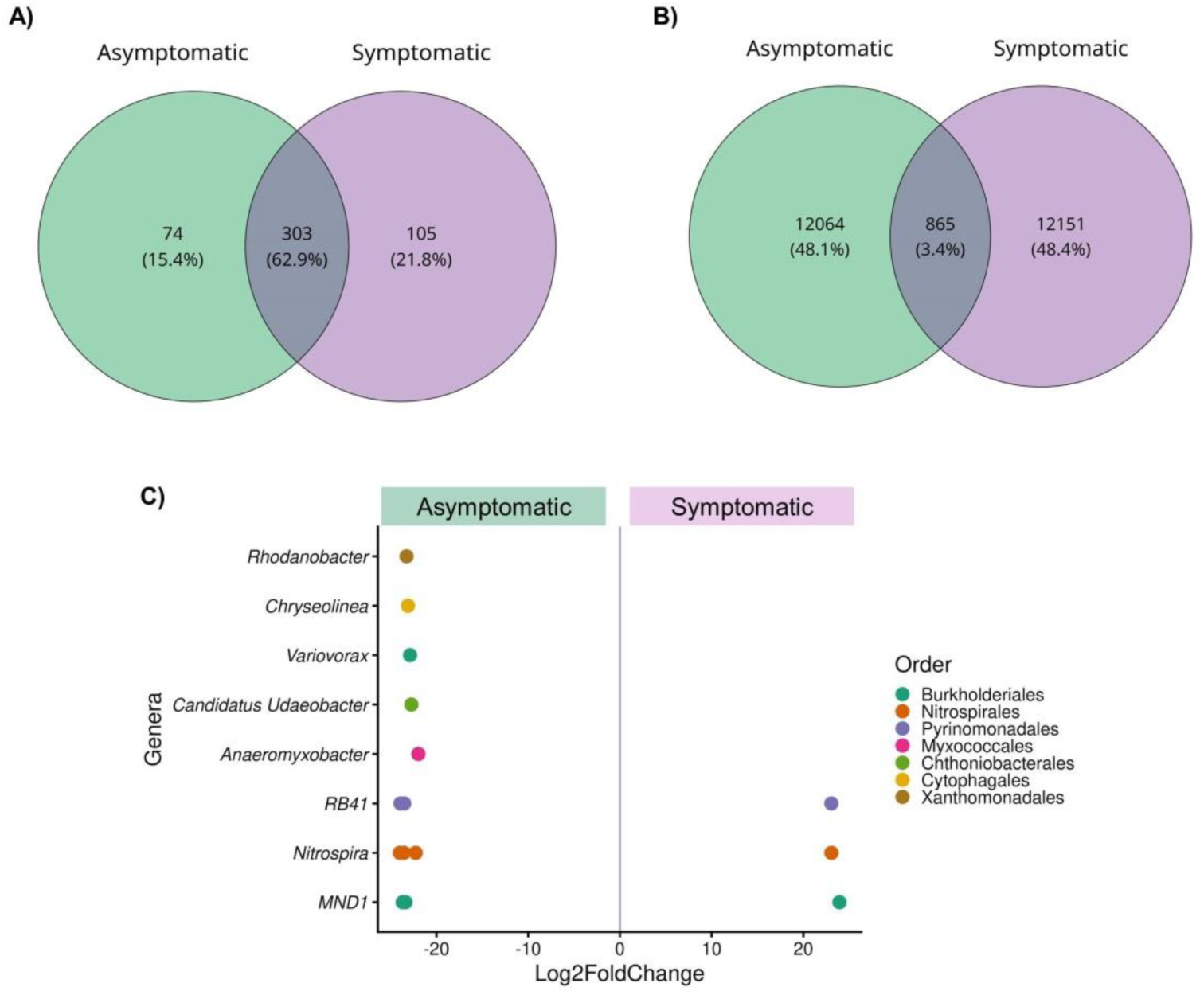
Shared and exclusive taxa in the rhizobacterial communities of symptomatic and PRR-symptomatic avocado trees. Venn diagrams of the number of unique and shared bacterial genera (A) and ASVs (B) between the two conditions. C) Differential abundance of bacterial ASVs identified at the genus level, associated with asymptomatic and PRR-symptomatic avocado trees according to the Wald test. Values of Log2FoldChange ≠ 0 and *P* < 0.05 are considered as differentially abundant in each condition.

### Functional diversity of the active rhizobacterial community of asymptomatic and PRR-symptomatic avocado trees

The composition of the active rhizosphere bacterial community was assessed using a comparative metatranscriptomic approach. Consistent with the results obtained from the metabarcoding data, alpha diversity indexes of the active bacterial communities did not vary between conditions (Supplementary Fig. 2A). Similarly, metatranscriptomic data obtained from asymptomatic and PRR-symptomatic rhizosphere soil samples did not display a clear separation in the Principal Component Analysis (PCA) (Supplementary Fig. 2B).

The differential expression analysis performed at the community level showed that 5,496 genes were up-regulated in the rhizosphere of asymptomatic trees whilst 5,891 genes were up-regulated in the PRR-symptomatic trees. A gene enrichment analysis was performed to cluster transcripts into GO terms that describe Biological Processes (BP) and Molecular Functions (MF). Interestingly, the bacterial community in the rhizosphere of PRR-symptomatic avocado trees showed an enrichment in transcripts of BP (37 *vs.* 1) and MF (23 *vs*. 1) of GO compared to asymptomatic trees. For asymptomatic avocado trees, enriched transcripts were affiliated to “de novo” protein folding (GO0006458) and protein folding chaperone (GO0044183) in the BP and MF categories of GO, respectively (Figure 3A and B). The most significantly enriched transcripts within BP in the PRR-symptomatic avocado trees were principally related to primary metabolism, such as phospholipid metabolic (GO0006644) and biosynthetic process (GO0008654), propionate metabolic process, methylcitrate process (GO0019679) and short-chain fatty acid metabolic process (GO0046459) (Figure 3A). In the MF category, the transcripts enriched in the PRR-symptomatic trees were related to the mRNA 3’-UTR binding (GO0003730), aconitase hydratase activity (GO0003994), 5S rRNA binding (GO0008097) and 2-methylisocitrate dehydratase activity (GO0047456) (Figure 3B).

**Figure 3.**
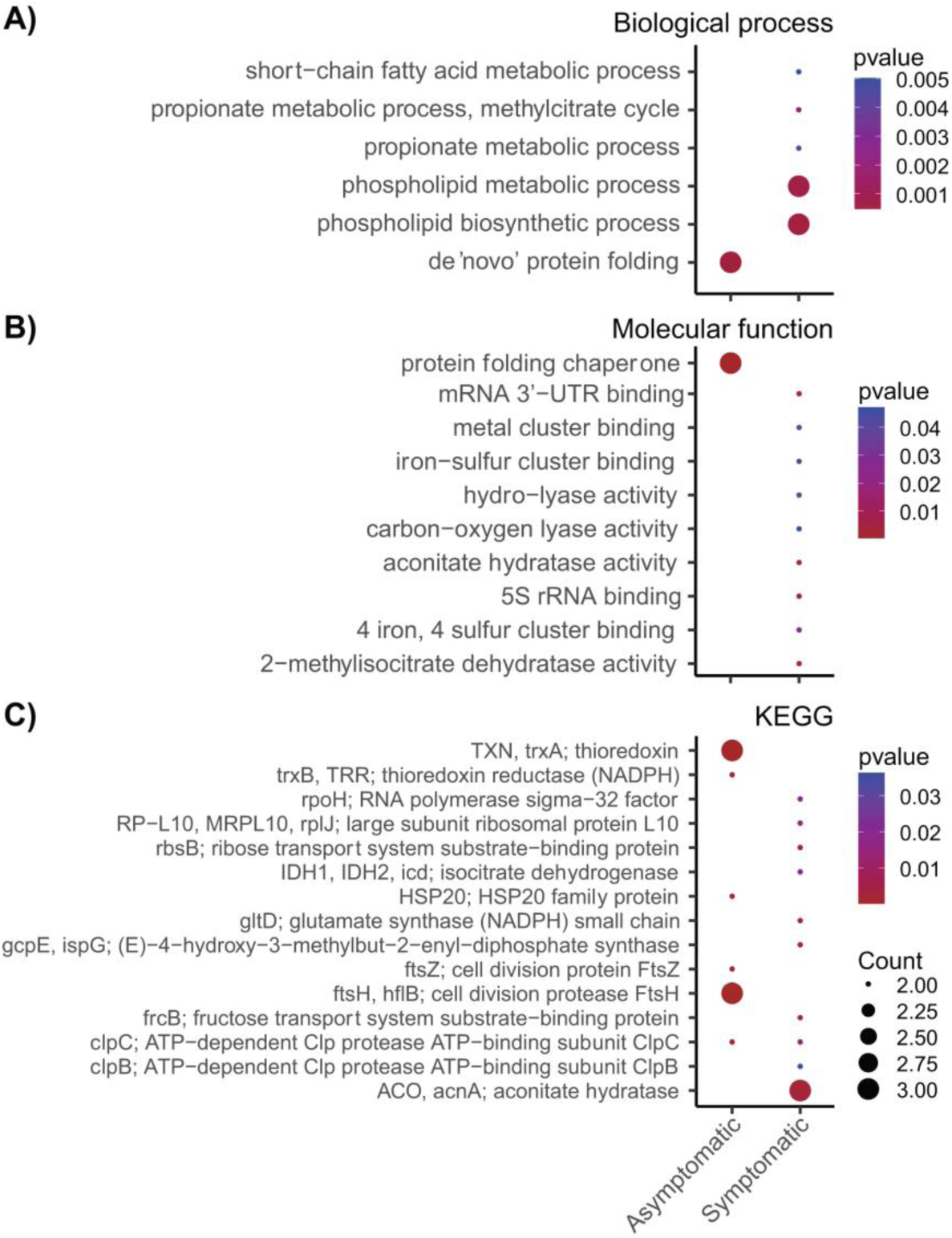
Gene Ontology (GO) and KEGG analysis of the active bacterial community in the rhizosphere of asymptomatic and PRR-symptomatic avocado trees. Functional enrichment analysis of asymptomatic and PRR-symptomatic metatranscriptomes based on GO Biological process enrichment (A) and Molecular function enrichment (B). C) KEGG Orthology (KO) enrichment of asymptomatic and PRR-symptomatic metatranscriptomes. Dot size represents the number of differentially expressed genes within the enriches of GO and KEGG and dot color represents the *P*-value (*P* < 0.05).

Similarly, KO analysis revealed more functions performed by the bacterial community in the rhizosphere of PRR-symptomatic trees than those from asymptomatic trees (33 *vs*. 15). The most enriched functions in asymptomatic avocado trees were the thioredoxin trxA (K03671: Protein families-genetic information processing), cell division protease FtsH (K03798: Protein families-metabolism) and HSP20 family protein (K13993: Genetic information processing). On the other hand, enriched functions in the PRR-symptomatic trees were related to aconitase hydratase (K01681: carbohydrate metabolism-citrate cycle (TCA), RNA polymerase sigma-32 factor (K03089: genetic information processing-transcription machinery), large subunit ribosomal protein L10 (K02864: genetic information processing-translation), ribose transport system substrate-binding protein (K10439: environmental information processing-membrane transport), isocitrate dehydrogenase (K00031: carbohydrate metabolism-TCA) and glutamate synthase (NADPH) small chain (K00266: energy metabolism-nitrogen metabolism) (Figure 3C).

### Composition and functional profile of active rhizobacteria

*Coxiella burnetti*, *Legionella pneumophila*, *Candidatus* Solibacter, *Gemmatimonas phototrophica*, *Gemmatimonas aurantiaca*, and *Streptomyces* sp. (Figure 4A) were amongst the most active rhizosphere bacterial taxa in both conditions. A differential expression analysis showed that *Mycoplasma haemofelis* (Log2FC = -7.26), *Anaeromyxobacter* sp. (Log2FC = -5.59), *Chroococcaceae* (Log2FC = -5.09), *Pilibacter termitis* (Log2FC = -5.09) and *Rickettsia rhipicephali* (Log2FC = -4.85) were the most significantly enriched bacterial genera in the rhizosphere of asymptomatic avocado trees. On the other hand, *Burkholderia* sp. (Log2FC = 7.26), *Hydrogenophaga* sp. (Log2FC = 7.18), *Rhodanobacter* sp. (Log2FC = 6.92), *Mycoplasma orale* (Log2FC = 6.81), *Modestobacter* sp. (Log2FC = 6.75), *Rhizobium* sp. (Log2FC = 6.65), and *Arthrobacter* sp. (Log2FC = 6.57) were significantly enriched in the rhizosphere of PRR-symptomatic avocado trees (Figure 4B).

**Figure 4.**
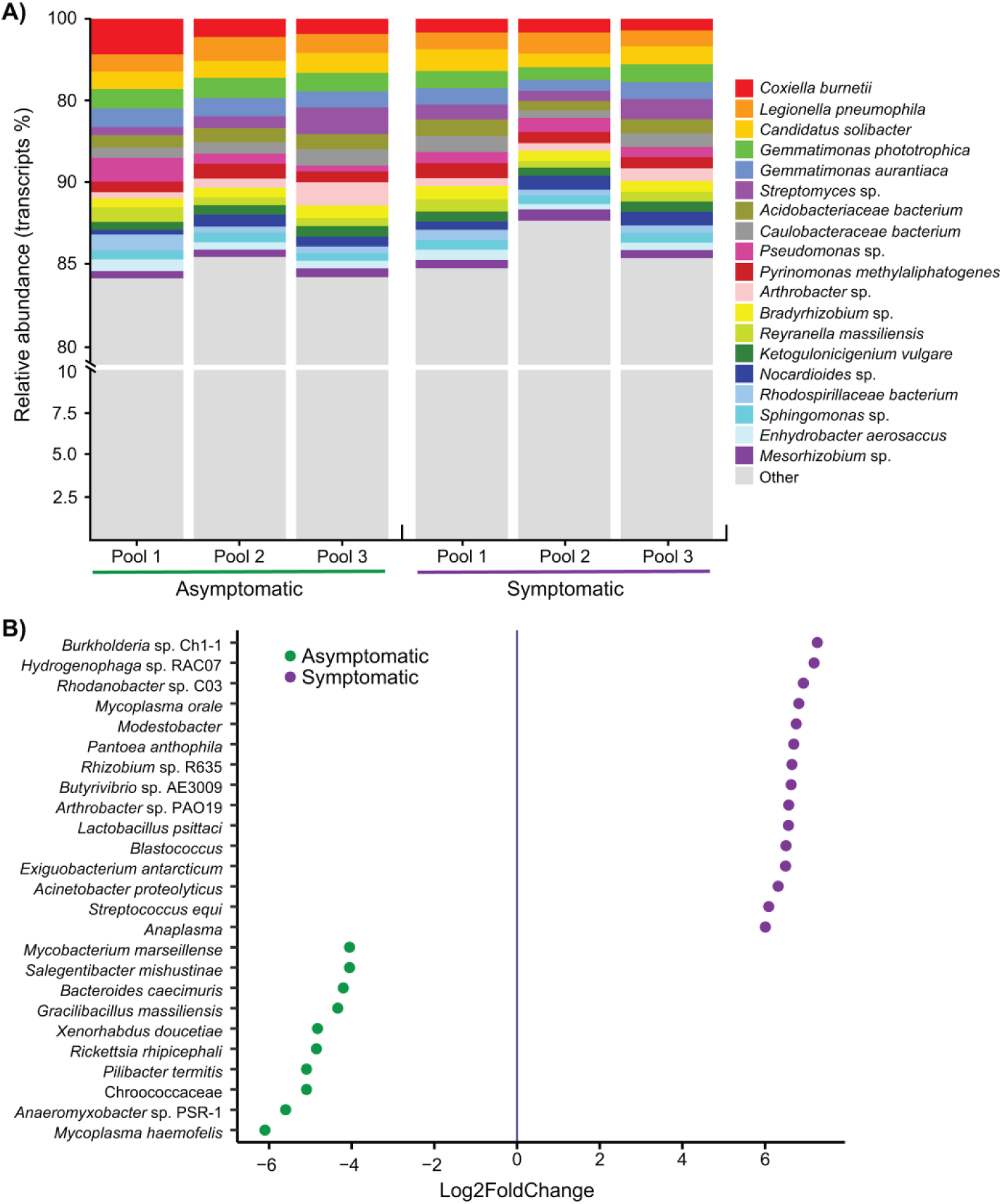
Taxonomic diversity of the active rhizobacterial community in the asymptomatic and PRR-symptomatic avocado trees. A) Relative abundance (%) of the most abundant bacterial species associated with asymptomatic and PRR-symptomatic avocado trees as retrieved by metatranscriptome taxonomy assignations. The others category represents bacterial species with a relative abundance < 1 %. B) Differential abundance of bacterial species associated with asymptomatic and PRR-symptomatic avocado trees according to the Wald test. Values of Log2FoldChange ≠ 0 and P < 0.05 are considered as differentially abundant in each condition.

The functions performed by the enriched genera in the PRR-symptomatic avocado trees were further analyzed. In accordance with results obtained in the functional profiling of the whole active community, *Burkholderia* sp., *Hydrogenophaga* sp., *Rhodanobacter* sp., *Rhizobium* sp. and *Arthrobacter* sp., the most active taxa from the PRR-symptomatic trees, expressed genes associated with stressful conditions and symbiotic processes, the biological functions pathogenesis and virulence, protein activation and nutrient acquisition, cell signaling, and bacterial and plant ion-homeostasis. Primary metabolism in these bacterial taxa was represented by genes encoding peptidases, transporters, kinases and hydrolases (Figure 5).

**Figure 5.**
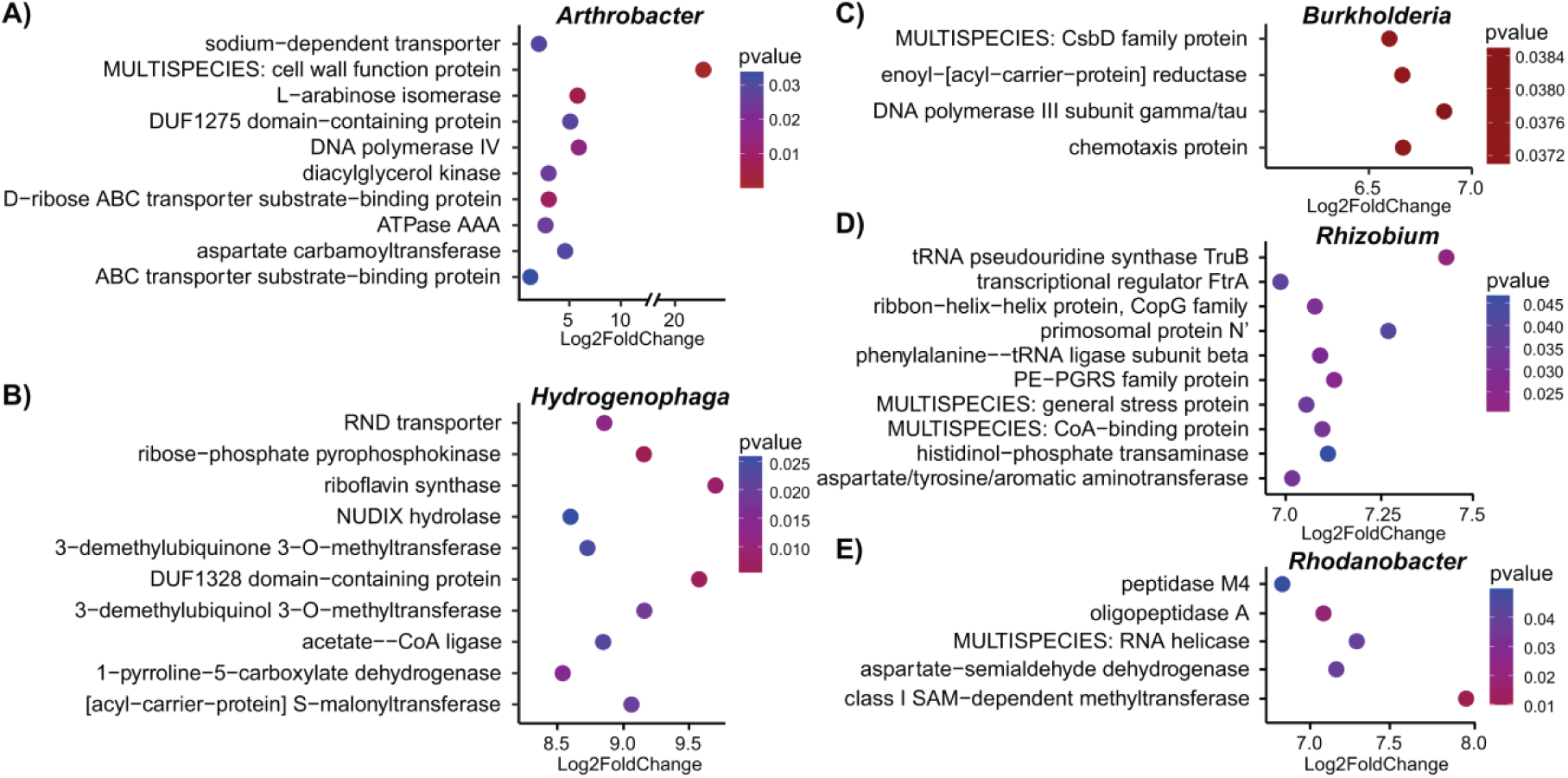
Functional profile of the most abundant and active bacterial genera in the rhizosphere of the PRR-symptomatic avocado trees. A) – E) Each panel enlists the top functions for each bacterium with Log2FoldChange values > 0 (PRR-symptomatic condition) and a *P*-value < 0.05.

## Discussion

*Phytophthora cinnamomi* is one of the most devastating pathogens infecting avocado trees. To better understand the impact of this pathogen on the avocado rhizosphere bacteriome, we combined metabarcoding with a metatranscriptomic approach to determine the compositional and functional shifts in avocado rhizosphere bacterial communities associated with PRR under field conditions. By investigating for the first time the impact of *P. cinnamomi* on the active bacteriome in the avocado rhizosphere, we showed that the enrichment of some bacterial genera in the PRR-symptomatic avocado trees may be related to the release of resources by root rot and that the active bacterial could be taking advantage of these released nutrients, as the metatranscriptomic analysis highlighted an activated carbohydrate metabolism in the rhizosphere of PRR-symptomatic trees. Moreover, we showed that enriched taxa in the rhizosphere of PRR-symptomatic trees were also associated with the expression of genes associated with a stress response, which suggests their active recruitment by the infected host. Collectively, our study aimed to contribute to the development of microbiome-based strategies that could be included into the integrated management of PRR in avocado.

The effect of soil-borne pathogens on the diversity of rhizosphere bacterial communities have been shown to vary, ranging from increases (Mendes et al., 2023; Yang et al., 2001) to decreases in rhizobacterial diversity in infected plants (Wei et al., 2018; Zhou et al., 2021), or with no effects (Hassani et al., 2023; Yin et al., 2021). Here, we found that *P. cinnamomi* did not affect alpha diversity metrics nor community structure (beta diversity) of the avocado rhizobacterial community, consistent with the findings of Solís-García et al. (2021) in Veracruz (Mexico) orchards. The lack of effect of *P. cinnamomi* on the structure and diversity of the avocado rhizosphere bacteriome could be explained by the distinct substrate preference by rhizobacteria and the pathogen, a necrotrophic oomycete. While bacteria may favor labile C sources (Wang & Kuzyakov, 2024), the oomycete displays an arsenal of glycoside hydrolases for the direct or indirect break-down of plant cell walls and plant cells, to feed on dead tissue (Bradley et al., 2022). As observed in other pathosystems, changes in diversity metrics may become more evident when pathogen infection induces an increased resource competition between the pathogen and the rhizosphere microbiome, e.g. between *Ralstonia solanacearum* and rhizobacteria in tobacco (Wang et al., 2022) or *Fusarium* and fungi in the sugarcane rhizosphere (Li et al., 2022). These findings suggest a potentially stronger effect of *P. cinnamomi* on rhizosphere fungal than bacterial communities and highlight the need for further study of inter-microbial nutrient competition in the rhizosphere of infected trees.

### Changes in the abundance of certain bacterial genera may be related to energy resources

Although *P. cinnamomi* did not significantly alter the structure and diversity of bacterial communities in the avocado rhizosphere, compositional shifts were detected in PRR-symptomatic trees compared to asymptomatic trees. Given that we did not find significant changes in soil chemical characteristics between the tree conditions, these shifts were likely induced by the presence of the pathogen.

Enriched bacterial taxa in the rhizosphere of asymptomatic avocado trees included *Variovorax*, *Rhodanobacter*, *Anaeromyxobacter*, *Candidatus* Udaeobacter and *Chryseolinea,* which may contribute to maintaining the health of their host. The *Variovorax* genus has been reported to promote plant growth, manipulating *Arabidopsis thaliana* hormones to balance the effects of a synthetic community of bacteria on root growth (Finkel et al., 2020) or enhancing wheat germination under salt stress conditions (Acuña et al., 2024). Moreover, Wang et al. (2022) found that *Variovorax* was more abundant in healthy roots of *Panax notoginseng* than in plants with symptoms of root rot and suggested that this genus could be a promising biological control agent against *P. notoginseng* root rot. Similarly, Pereira et al. (2025) reported that *Variovorax paradoxus* exhibited *in vitro* antagonistic activity against *F. oxysporum*. On the other hand, the enrichment of *Rhodanobacter* in the ginseng rhizosphere was shown to increase its resistance to disease, improving crop yield and quality (Li et al., 2024). Moreover, novel strains of *Rhodanobacter* have displayed antagonistic activities *in vitro* against the fungal pathogens *Cylindrocladium spathiphylli* (De Clercq et al., 2006) and *Fusarium solani* (Huo et al., 2018). Regarding *Chryseolinea* and its interactions with plant pathogens, results from the literature are contrasting. Ou et al. (2019) found that *Chryseolinea* may play a role against *Fusarium oxysporum* establishment in disease suppressive soils, while Tang et al. (2023) described a positive correlation of this taxa with *Fusarium* wilt incidence; further studies are thus necessary to better understand the possible role of *Chryseolinea* in mitigating *P. cinnamomi* infection. Overall, these reports suggest that the presence of these bacteria in the rhizosphere of asymptomatic avocado trees could be associated with the health status of the plants.

The significant increase in ASVs from the genera MND1, RB41 and *Nitrospira* in both PRR-symptomatic and asymptomatic rhizosphere soil samples could respond to different mechanisms between conditions. Wei et al. (2024) reported the enrichment of these three genera in rhizosphere soils from healthy tobacco plants compared to plants infected by *R. solanacearum* and suggested that these taxa may be related to plant-root symbiosis and nutrient acquisition, nitrogen cycling and organic matter decomposition. Moreover, they found that RB41 and MND1 positively correlated with seven active metabolites involved in the synthesis of antibiotics and alkaloids (Wei et al., 2024), which may indicate their role in disease suppression. MND1 has also been described as being more abundant in the rhizosphere of healthy plants compared to plants infected by *R. solanacearum* (Wen et al., 2020). On the other hand, members of the genus *Nitrospira* have been described as complete ammonia oxidizers in forest, grassland and agricultural soils (Hu et al., 2021), thus playing an important role in soil nitrification. *Nitrospira* was also found to be negatively related to disease incidence in tobacco, putatively due to its contribution to nitrate production and plant disease resistance (Li et al., 2025). Collectively, these results indicate that the enrichment of MND1, RB41 and *Nitrospira* in the rhizosphere of PRR-symptomatic trees may be due 1) the release of resources due to root rot and 2) their recruitment as part of the plant cry-for-help, as important contributors to plant health and defense response (Mur et al., 2016; Sun et al., 2021). Further functional analyses were thus required to corroborate the potential beneficial effect of these bacterial taxa to disease mitigation and elucidate whether their enrichment results from an increase in available carbohydrates in the rhizosphere of infected trees or from a plant defense response (Alfaro-García et al., 2025).

### Rhizobacteria may be responding to the release of resources by *Phytophthora cinnamomi* root-rot

The functional enrichment analysis based on COG and KEGG databases showed that the active bacterial community changed depending on health status of the host tree. We found more enriched pathways in the rhizosphere of PRR-symptomatic trees than in that of asymptomatic avocado trees. The enriched metabolic routes in infected trees were generally related to carbohydrates metabolism, nitrogen metabolism and to post-transcriptional and translation regulation.

The reiterative enrichment of the aconitate hydratase pathway in infected trees in both COG and KEGG annotations suggests that carbohydrate metabolism is stimulated in the rhizobacterial communities associated with PRR-symptomatic trees. This enzyme participates in the tricarboxylic acid (TCA) cycle, catalyzing the reversible hydration of cis-aconitate to produce citrate or isocitrate. Additionally, the enrichment of the aconitate hydratase in symptomatic trees combined with that of 2-methylisocitrate dehydratase confirms that the 2-MC cycle is occurring too. Both the TCA and 2-MC cycles are coupled, since they share enzymes such as the aconitate hydratase and the 2-MC cycle provides substrates for the TCA cycle. The significant enrichment of the 4iron-4sulfur cluster (COG), another key player in carbon metabolism, in the rhizosphere of infected trees, indicates that active rhizosphere bacteria in symptomatic avocado trees are likely taking advantage of the C resources released by root rot. These findings are consistent with those by Shu et al. (2019), who detected with metagenomics an increase in genes related to carbohydrate metabolism in the rhizosphere of avocado trees infected by *P. cinnamomi* and suggested that the presence of the pathogen stimulates the primary metabolism of the rhizosphere bacterial community.

On the other hand, transcripts related to maintenance of the cellular proteostasis network, which allows bacteria to acclimatize to abiotic stress, were enriched in the asymptomatic condition. The enrichment of chaperone protein families such as Hsp20 and protein folding chaperone confirms the role of the rhizobiome in providing stress tolerance to the host plant; the heat shock protein Hsp20 has been reported to increase following exposure to stress, including temperature changes (Xue et al., 2019) o desiccation resistance (León-Sobrino et al., 2024). Thioredoxin systems (trxA and trxB) act as a sensor favoring stress adaptation, principally to reactive oxygen species (ROS), O_2_ and reactive N species. These protein systems control the modifications of oxidized proteins (thiol group), ensuring protein repair and maintaining the thiol homeostasis in the cell (Anjou et al., 2024). FtsH and Clp proteases belong to the Hsp100 family proteases. Clp proteins are involved in stress response, sporulation, swarming, motility and biofilm formation, whereas FtsH metalloprotease is involved in sporulation initiation, biofilm formation, cell envelope stress, heat shock and is directly involved in protein quality control through the degradation of misfolded or damaged proteins (Harwood & Kikuchi, 2021). Overall, the depletion of these transcripts in the PRR-symptomatic avocado trees could indicate a loss of resilience in the event of additional stressors, such as drought or nutrient scarcity.

### The dominant active bacteria in symptomatic avocado trees are associated with plant growth-promotion

The active bacterial communities in the rhizosphere of PRR-symptomatic avocado trees were dominated by *Burkholderia*, *Hydrogenophaga*, *Rhodanobacter*, *Modestobacter*, *Rhizobium* and *Arthrobacter*. Most of these taxa have been associated with plant growth promotion and disease suppression, which suggests they could have been recruited by the plant upon infection to assist its defense response, although we cannot discard the possibility that these taxa were attracted by the nutrient release following root necrosis. For example, *Rhodanobacter*, which was enriched in the rhizosphere of asymptomatic avocado trees (metabarcoding), was detected as more active in the rhizosphere of infected trees (metatranscriptomics), with an increased expression of peptidase M4 and oligopeptidase (Hasan et al., 2021). This may indicate an increase in resource acquisition in a nutrient-rich condition.

However, some of the enriched transcripts in these bacterial taxa point at a possible recruitment by the infected plant. For example, *Burkholderia* and *Arthrobacter* strains have been isolated from the sugarcane rhizosphere and displayed antagonistic activity against *Fusarium commune* GXUF-3, suppressing its mycelial growth and spore germination (Li et al., 2022). In particular, the plant growth-promoting activity exhibited by *Burkholderia* strains was attributed to their N-fixing capacity. One of the differentially expressed transcripts associated to *Burkholderia* in the present study was chemotaxis protein. Bacteria use chemotaxis to sense changes in chemical concentration gradients in the environment, allowing them to search for food, avoid toxins and respond to changing environmental conditions (Wadhams & Armitage, 2004). *Burkholderia* spp. have been reported to display a positive chemotactic response to the root exudates of *Panax notoginseng* in autotoxic ginsenoside stress, which allowed them to enhance their root colonization and protect the plant from root rot disease (Deng et al., 2023). These findings support the cry-for-help hypothesis behind the enrichment of *Burkholderia* in PRR-infected avocado trees.

Similarly, *Arthrobacter* has also been reported for its anti-oomycete and plant growth-promoting activities in several plant species (Méndez-Bravo et al., 2018; Town et al., 2016; Velázquez-Becerra et al., 2013) and may have also been recruited by the infected plant as part of a cry-for-help strategy. In the rhizosphere of PRR-symptomatic avocado trees, *Arthrobacter* was actively transporting substrates from the surrounding environment, as transcripts related to ABC transporters (D-ribose, substrate-binding protein) were differentially expressed for this genus. ABC transporters are involved in the import and export of a wide range of substrates, such as sugars, amino acids, ions and complex organic molecules (Tanaka et al., 2018), supporting the hypothesis that *Arthrobacter* may have benefited from the release of carbohydrates following root rot (Alfaro-García et al., 2025). However, D-ribose has also been reported, among other carbohydrates, to stimulate the growth and diversity of tomato rhizobacteria, resulting in a reduction of the pathogen *R. solanacearum* abundance (Wen et al., 2023). D-ribose production occurs through the pentose phosphate pathway (PPP), which has an important role in maintaining redox homeostasis and activating stress responsive gene expression under oxidative stress (Gupta & Gupta, 2021). As H_2_O_2_ has been described to increase in the interaction of avocado-*P. cinnamomi* (García-Pineda et al., 2010), the activation of PPP could help rhizobacteria to deal with the ROS produced by the avocado during infection.

## Conclusions and final considerations

Our study confirmed the dysbiosis caused by *P. cinnamomi* on the avocado rhizobacterial community, at the compositional and functional levels. The active bacterial community associated with PRR-symptomatic avocado trees displayed an induced gene repertoire ranging from an enhanced carbohydrate metabolism to the expression of genes related to stress response, cell signaling, ion-homeostasis, virulence factors, and symbiotic interactions with plants. Our findings highlight the importance of combining compositional and functional microbiome analyses to distinguish between microbial taxa recruited by the plant as part of a cry-for-help strategy and those merely attracted by the release of labile C compounds following root rot. Future studies will aim at complementing our understanding of the avocado microbiome dysbiosis induced by *P. cinnamomi*, by investigating the effect of the pathogen on rhizosphere fungal communities and on root exudation patterns through metabolomics. Collectively, our multi-omics study of the avocado-*P. cinnamomi* pathosystem contributes to establish the bases for future microbiome manipulation strategies and for the build-up of disease-suppressive soils that could help mitigate the impact of this important soil-borne pathogen.

## Acknowledgements

This study was funded by project FOSEC-SEP-CONAHCYT (SECIHTI), grant number A1-S-30794 and by the CONAHCYT(SECIHTI)-PRONACES-PPE grant (project “PERSEA”, 322772). R.A.-G. was granted with a SECIHTI PhD scholarship (CVU 932217). We are grateful to Alejandra Mondragón-Flores (INIFAP) and Sylvia Fernández-Pavía (UMSNH) for their help with orchard selection and to Ing. Victor for allowing us the access to the orchard. We also thank Daniel Sánchez-Hernández and Alejandra Mondragón-Flores for their help with sampling and Alejandro Soto-Plancarte for his help with field work and confirmation of *P. cinnamomi* presence. Special thanks to Laura Piñon and Eduardo Thome for their technical assistance in the sequencing libraries construction and to Diego Isla López for his technical assistance with computational tools at Cluster-LANASE, ENES-Morelia, UNAM. We also thank Sandra Rocha and colleagues for their work with the soil chemical analysis.

## Notes

### Competing Interest Statement

The authors have declared no competing interest.

