## Supplementary Figures for "*Phytophthora* root rot induces compositional and functional changes in avocado rhizosphere bacterial communities"

[3] Laboratorio de Ciencias Agrogenómicas and Laboratorio Nacional PlanTECC, Universidad Nacional Autónoma de México. Blvd. UNAM 2011, Predio El Saucillo y Comunidad Los Tepetates, El Potrero. 37684 León, México.

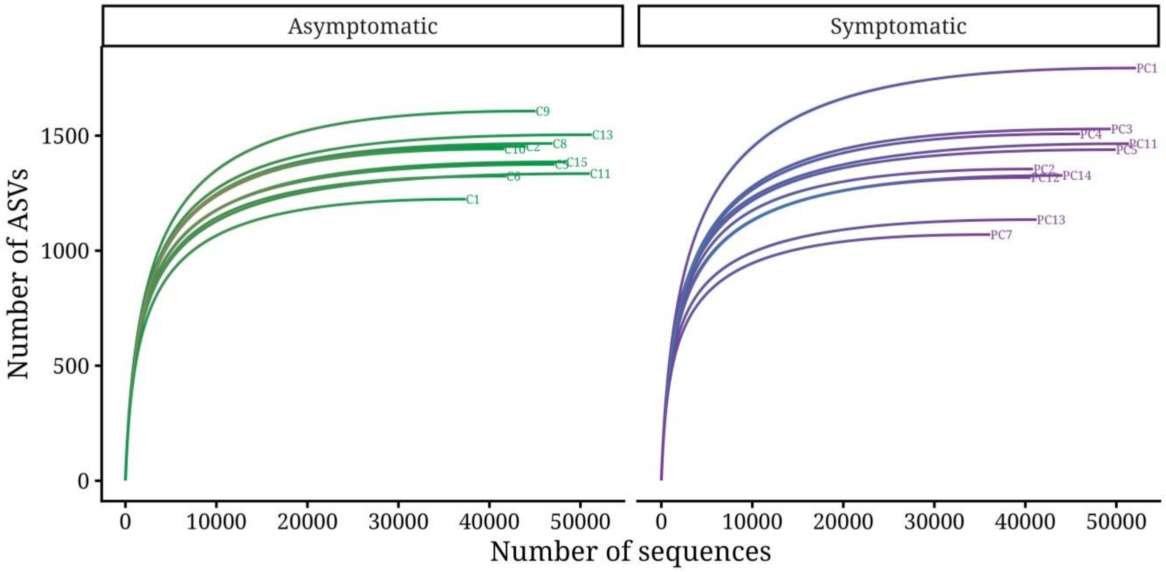


7

**Supplementary Figure 1. Rarefaction curves of the observed bacterial ASVs associated with asymptomatic and PRR-symptomatic avocado trees.**

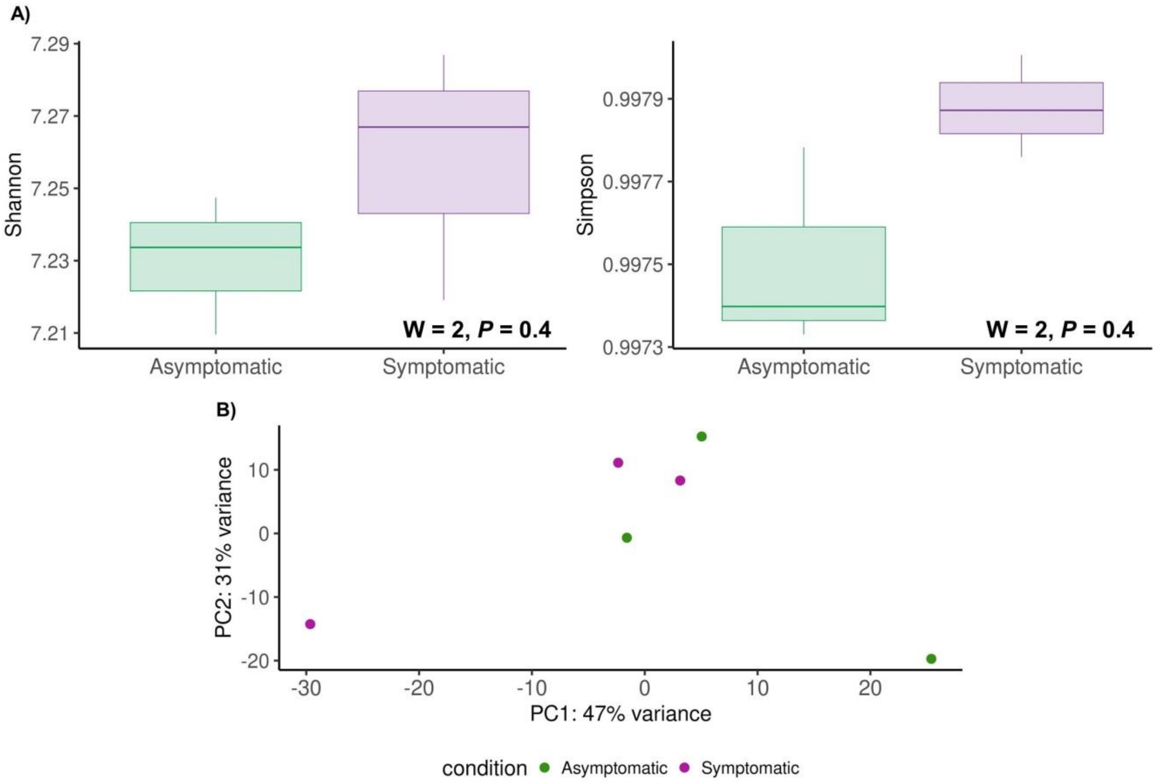


**Supplementary Figure 2. Diversity and structure of the active bacterial community in the rhizosphere of asymptomatic and PRR-symptomatic avocado trees.** A) Shannon and Simpson indices of active rhizobacterial communities associated with asymptomatic and PRR-symptomatic avocado trees. The W and *P* values were calculated with the Mann-Whitney-Wilcoxon test. B) Principal component analysis (PCA) showing the functional structure of asymptomatic and PRR-symptomatic avocado trees retrieved from expression data across all the metatranscriptome.
